# Faster diffusive dynamics of histone-like nucleoid structuring proteins in live bacteria caused by silver ions

**DOI:** 10.1101/776229

**Authors:** Asmaa A. Sadoon, Prabhat Khadka, Jack Freeland, Ravi Kumar Gundampati, Ryan Manso, Mason Ruiz, Venkata R. Krishnamurthi, Suresh Kumar Thallapuranam, Jingyi Chen, Yong Wang

## Abstract

The antimicrobial activity and mechanism of silver ions (Ag^+^) have gained broad attention in recent years. However, dynamic studies are rare in this field. Here, we report our measurement of the effects of Ag^+^ ions on the dynamics of histone-like nucleoid structuring (H-NS) proteins in live bacteria using single-particle tracking photoactivated localization microscopy (sptPALM). It was found that treating the bacteria with Ag^+^ ions led to faster diffusive dynamics of H-NS proteins. Several techniques were used to understand the mechanism of the observed faster dynamics. Electrophoretic mobility shift assay on purified H-NS proteins indicated that Ag^+^ ions weaken the binding between H-NS proteins and DNA. Isothermal titration calorimetry confirmed that DNA and Ag^+^ ions interact directly. Our recently developed sensing method based on bent DNA suggested that Ag^+^ ions caused dehybridization of double-stranded DNA (i.e., dissociation into single strands). These evidences led us to a plausible mechanism for the observed faster dynamics of H-NS proteins in live bacteria when subjected to Ag^+^ ions: Ag^+^-induced DNA dehybridization weakens the binding between H-NS proteins and DNA. This work highlighted the importance of dynamic study of single proteins in the live cells for understanding the functions of antimicrobial agents to the bacteria.

## Introduction

Due to the rise of antibiotic resistance of bacteria ^[1]^, alternatives to traditional antibiotics have been attracting broad interest and attention toward fighting against bacterial infections ^[2,3]^. A promising candidate among the available alternatives is silver (Ag), which has long been known and used as an antimicrobial agent, dating back as far as ancient Greece, the Roman Empire, and Egypt ^[4]^. In the past decades, the potent antimicrobial properties of Ag have been revisited in various forms such as ions, surfaces, and nanoparticles, and exciting progresses have been made ^[5–16]^. For example, it has been reported that damages in bacteria caused by Ag are multimodal, including DNA condensation and damage, free radical generation (ROS), and loss of ATP production ^[17–22]^. However, the exact mechanism underlying the antimicrobial activity of Ag remains not fully understood ^[17]^. Most of the existing literature relied on the traditional bioassays for the mechanistic studies. Little effort has been made in studying the molecular dynamics inside the bacteria; therefore, temporal resolution for understanding the damages in bacteria caused by Ag is still missing ^[17]^.

In this work, we used super-resolution fluorescence microscopy ^[23–26]^ in combination with single-particle tracking ^[27–33]^ to investigate and understand the effects of Ag^+^ ions on the dynamic diffusion of individual proteins at the molecular level in live *E. coli* bacteria. The protein in this study is the histone-like nucleoid-structuring (H-NS) protein ^[34]^, which was chosen for the following three reasons. First, the H-NS protein is an essential protein in *E. coli*, as determined by Gerdes et al. ^[35]^, and serves as a universal negative regulator, regulating (mostly negatively) ~5% of the bacterial genome ^[34,36]^ (Fig. 1a). Second, the H-NS protein is tightly associated with various biological processes in the bacteria that respond to damages due to Ag, such as modulating the synthesis and stability of RpoS – a central protein/regulator for general stress responses ^[37–39]^, compacting DNA and cause DNA condensation ^[40,41]^, modulating the production of deoxyribonucleotides and synthesis of DNA ^[42]^, and enhancing the cellular defenses against ROS ^[37]^. Third, nanoscale spatial reorganization of H-NS proteins has been previously observed to form denser and larger clusters in bacteria subjected to Ag-treatment ^[43]^.

**Figure 1.**
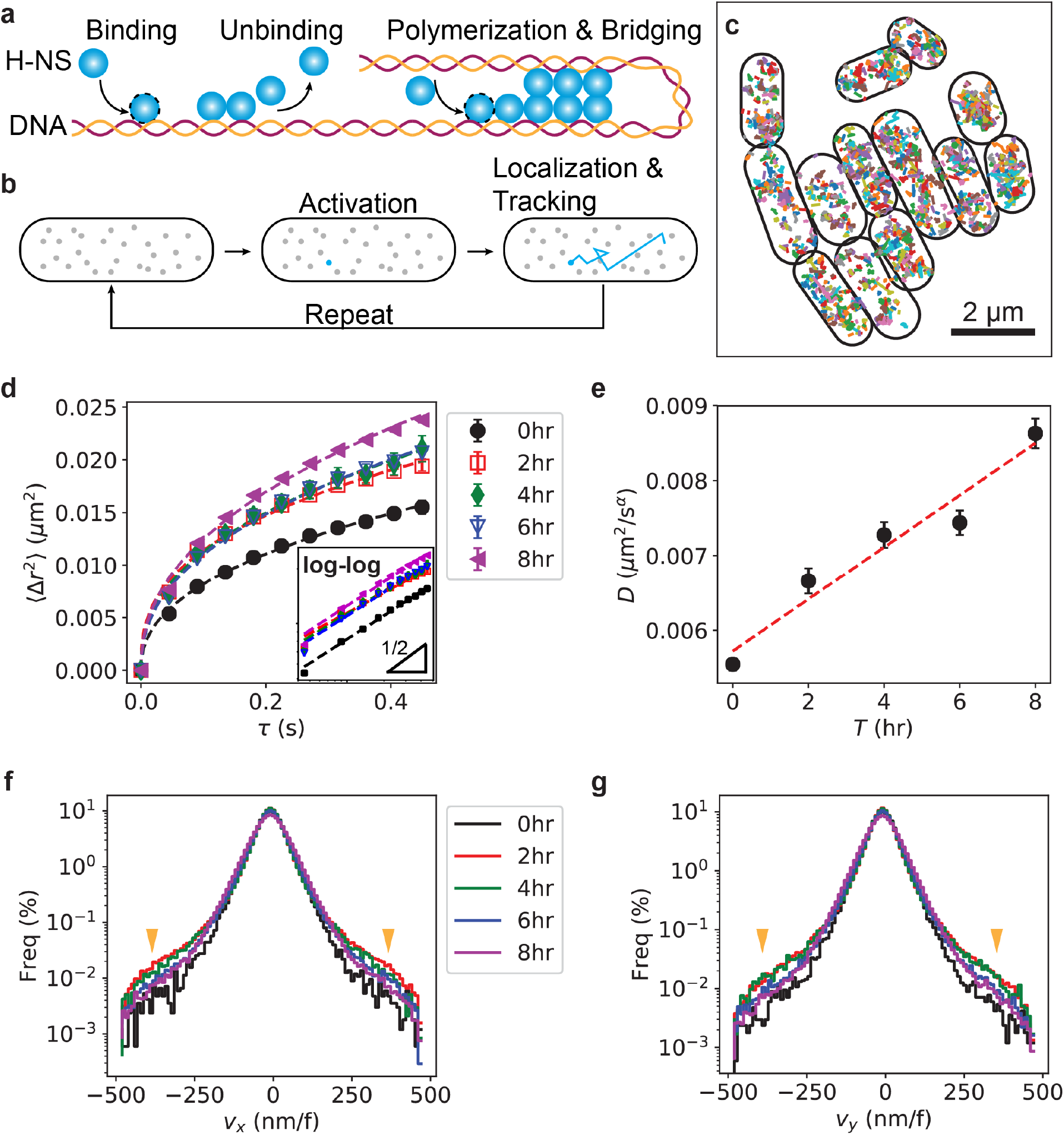
Faster dynamics of H-NS proteins in live bacteria caused by Ag^+^ ions. (a) Illustration of H-NS proteins’ key activities. H-NS is a DNA-binding protein, consisting of a DNA-binding domain, a linker, and an oligomerization domain, which allows H-NS to form polymers and DNA bridging. (b) Illustration of sptPALM for tracking H-NS proteins in live *E. coli*. (c) Examples of trajectories of H-NS proteins in individual *E. coli* in a region of interest (ROI) of 8×8 µm^2^. (d) Ensemble mean-square-displacement (eMSD) of H-NS proteins in live *E. coli* bacteria in the absence of Ag^+^ ions (closed black circles), or in bacteria treated by 10 µM Ag^+^ ions for 2 hours (open red squares), 4 hours (closed green diamonds), 6 hours (open blue triangles), and 8 hours (closed magenta triangles). Symbols stand for averages of eMSD data over *E. coli* cells (the number of cells ranges from 158 to 678). Error bars represent the standard errors of the means (SEM). Dashed curves are fittings using 〈Δ*r*^2^〉 = 4*Dτ*^*α*^ where *D* is the generalized diffusion coefficient and *α* is the anomalous scaling exponent. Inset: the same data plotted in log-log scale, with a slope of ½ indicated. (e) Dependence of the generalized diffusion coefficient *D* on the treatment time *T*. Error bars represent fitting errors. (f, g) Log-linear distributions of instantaneous velocities (C: *v*_*x*_; D: *v*_*y*_) of H-NS proteins in the absence (black) and presence of 10 µM Ag^+^ ions (colored lines) for different durations. The orange arrows highlight the increase of the probability of higher velocities after Ag^+^-treatment.

Using single-particle tracking photoactivated localization microscopy (sptPALM, Fig. 1b) ^[27,28]^ with a spatial resolution of 20 nm and a temporal resolution of 45 ms, we observed that treating the bacteria with Ag^+^ ions led to faster dynamics of H-NS proteins. While the motion of H-NS proteins in live bacteria after Ag^+^-treatment remains sub-diffusive, the generalized diffusion coefficient of H-NS proteins increased when the bacteria were exposed to Ag^+^ ions for longer period of time. Analyzing the step sizes of the diffusion of H-NS proteins showed that treatment with Ag^+^ ions caused higher instantaneous velocities of this proteins. To understand the mechanism of the observed faster dynamics of H-NS, electrophoretic mobility shift assay was performed with purified H-NS proteins and double-stranded DNA. It was found that Ag^+^ ions weaken the binding between H-NS proteins and the DNA. Measurements with isothermal titration calorimetry suggested that Ag^+^ ions directly interact with DNA. Furthermore, we examined the effect of the DNA-Ag interaction, using our recently developed sensors based on bent DNA molecules, and found that Ag^+^ ions induced dehybridization of double-stranded DNA (i.e., dissociation into single strands). Based on these evidences, we provided a plausible mechanism for the faster dynamics of H-NS proteins in live Ag-treated bacteria. This work presents the first dynamic study at the molecular level – with both high spatial resolution and temporal resolution – on the antimicrobial mechanism of Ag^+^ ions. It is an important milestone for furthering our understanding the interactions of Ag to bacteria in motion quantitatively.

## Results

### Faster diffusive dynamics of H-NS proteins in live bacteria caused by Ag^+^ ions

Single-particle tracking photoactivated localization microscopy (sptPALM) was used to monitor the dynamic diffusion of H-NS proteins in live *E. coli*, as illustrated in Fig. 1b and described in “Methods” ^[32]^. Examples of diffusive trajectories of H-NS proteins in individual untreated bacteria in an area of 8×8 µm^2^ were shown in Fig. 1c. The lengths of the H-NS trajectories are not apparently affected by Ag^+^-treatment, with an average length of ~3 frames and maximum lengths above 100 frames (Fig. S1). From the trajectories, the ensemble mean-square-displacements (eMSD), 〈Δ*r*^2^(*τ*)〉 = 〈(***r***(*t* + *τ*) − ***r***(*t*))^2^〉, were calculated as shown in Fig. 1d, where the error bars (smaller than the symbols in some cases) represented the standard errors of the means (SEM) ^[32]^. It is noted that short trajectories were not removed in the calculations of eMSD calculations for two reasons. First, the determination of the cutoff lengths for “short” trajectory is arbitrary and prone to human bias. Second, although short trajectories affect the accuracy for calculating the individual MSD (iMSD) from single trajectories for single molecules, the eMSD is expected to be unaffected due to the ensemble averaging.

We observed that the eMSD curves for H-NS proteins in the treated bacteria were higher compared to the untreated ones (Fig. 1d), suggesting that the diffusive dynamics of H-NS proteins in live bacteria became faster after treatment with Ag^+^ ions, as the eMSD is positively related to the diffusion coefficient, 〈Δ*r*^2^(*τ*)〉 = 4*Dτ*^*α*^ in 2D, where *D* is the generalized diffusion coefficient and *α* is the anomalous scaling exponent. This observation can be confirmed in the log-log plots of the eMSD curves (inset of Fig. 1d), where the y-intercepts represent the diffusion coefficients and the slopes indicate the anomalous scaling exponent *α* as log〈Δ*r*^2^〉 = log 4*D* + *α* ⋅ log *τ*. The eMSD curves were shifted upwards for treated bacteria, showing higher y-intercepts and confirming that the generalized diffusion coefficients of the H-NS proteins were higher in bacteria treated with Ag^+^ ions. In contrast, we observed that the anomalous scaling exponent *α* was not significantly affected by the treatment with 10 µM Ag^+^ ions, as the slopes of eMSD curves in the log-log scale were similar for the untreated and Ag^+^-treated bacteria (~0.5, inset of Fig. 1d).

To quantify the faster diffusive dynamics caused by Ag^+^ ions, we fitted the eMSD data using 〈Δ*r*^2^〉 = 4*Dτ*^*α*^ and obtained the fitted generalized diffusion coefficients for the untreated bacteria and bacteria treated with 10 µM Ag^+^ ions for 2, 4, 6, and 8 hr. It is noted that our previous results showed that treating *E. coli* with Ag^+^ ions at 10 µM extended the lag time of the bacterial culture, while the growth rate remained the same ^[43,44]^. As shown in Fig. 1e, the diffusion coefficient steadily increased as the treatment time increased. Fitting the data with a linear line suggested that the diffusion coefficient increased [(3.5 ± 0.5) × 10^AB^ µm^2^/s per hour]. After 8 hr, a significant increase of 56% in the diffusion coefficient was observed. We point out that, as the concentration of the added Ag^+^ ions (10 µM) is too low compared to the ionic strength of the medium (~100 mM Na^+^/K^+^ and 2 mM Mg^2+^), simple ionic effects cannot result in the observed increase of the diffusion coefficient.

To further examine the faster dynamics of H-NS proteins in live bacteria, we estimated the instantaneous velocities of the proteins from the trajectories using 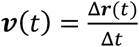, where Δ**r** is the displacement and Δ*t* is the time interval between adjacent data points in the trajectories. Note that, as we used a memory of 1 frame (Mem = 1) when linking trajectories, Δ*t* could be either 45 ms (time interval between adjacent frames) or 90 ms. The histograms of the x- and y-components of the instantaneous velocities (*v*_*x*_ and *v*_*y*_) are shown in Fig. 1f and 1g. Similar to our previous observations ^[32]^, the distributions of the velocities deviated from the Gaussian distribution at higher velocities, confirming the abnormality of dynamic diffusion of H-NS proteins in live bacteria. Further, we observed that the fraction of larger velocities was higher after subjecting bacteria to Ag^+^ ions compared with the untreated bacteria (Figs. 1f and 1g). This observation suggested that, upon Ag^+^-treatment, the probability of H-NS proteins travelling with larger velocities increased. Our previous study suggested that the deviation of the distribution of the instantaneous velocities from the Gaussian distribution is likely due to H-NS proteins’ binding/unbinding on DNA ^[32]^. Therefore, the observed changes in the velocity distributions suggested that treating live bacteria with Ag^+^ ions probably affected the binding of H-NS proteins on DNA.

Two parameters are important for identifying and linking trajectories of molecules: the maximum displacement that a particle can move between frames (Maxdisp) and the maximum number of frames during which a particle can vanish (the memory or Mem) ^[27–33]^. To assess the robustness of the observations, we used different values for these two parameters (Maxdisp ranging from 400 to 560 nm, and Mem ranging from 0 to 2). We observed that results with the different parameters (Fig. S2, S3 and S4) were similar to the ones using Maxdisp = 480 nm and Mem = 1 (shown in Fig. 1). Therefore, the observation of the faster diffusion of H-NS proteins caused by Ag^+^ ions is robust.

### Weaker binding of H-NS proteins on DNA due to Ag^+^ ions

The observed faster dynamics of H-NS proteins due to Ag^+^-treatment led to a hypothesis that Ag^+^ ions promoted dissociation of H-NS proteins from the chromosomal DNA of the bacteria. To test this hypothesis, we expressed and purified H-NS proteins (with a purity of ~50% based on quantification using PAGE gel electrophoresis) and performed *in vitro* electrophoretic mobility shift assay (EMSA) ^[45]^. Representative gel images showing the bands of unbound DNA are presented in Fig. 2a. In each gel, the concentration of DNA was fixed, and the concentration of H-NS proteins increased from 0 to 433 µM (from left to right). The concentration of Ag^+^ ions changed in different gels, ranging from 0 to 1 mM (Fig. 2a). In the absence of Ag^+^ ions (0 mM), the amount of unbound DNA decreased steadily as the concentration of H-NS proteins increased, indicating the binding of H-NS proteins to the double-stranded DNA. The bands for the unbound DNA almost disappeared for the last two lanes (corresponding to 346 and 433 µM, respectively). In contrast, the bands of unbound DNA at the same concentration of H-NS proteins showed higher intensities in the presence of Ag^+^ ions (Fig. 2a).

**Figure 2.**
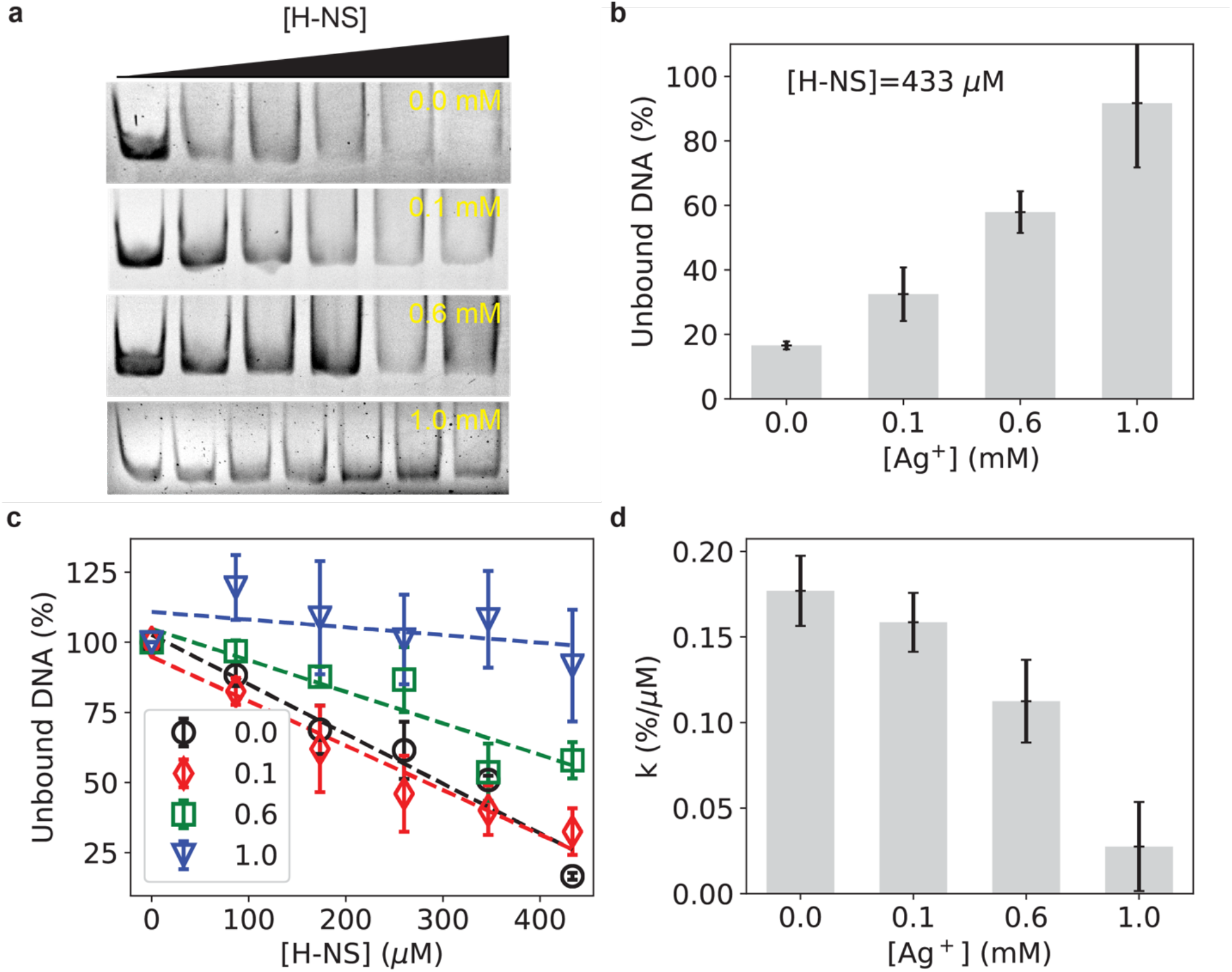
Electrophoretic mobility shift assay (EMSA) for measuring the binding between H-NS proteins and double-stranded DNA in the absence and presence of Ag^+^ ions. (a) Examples of EMSA gels for H-NS proteins in the absence (0 mM) and presence of Ag+ ions (0.1 mM, 0.6 mM, and 1.0 mM). In each gel, the concentrations of DNA and Ag^+^ ions were fixed, while the concentration of H-NS protein increased linearly from left to right of the gel (0, 87 µM, 173 µM, …, 433 µM). (b) Dependence of the percentage of unbound DNA on the concentration of Ag^+^ ions in the presence of 433 µM H-NS proteins (last lanes in panel a). (c) Dependence of the normalized amount of unbound DNA on the concentration of H-NS proteins in the absence (black circles) or in the presence of Ag^+^ ions (red diamonds: 0.1 mM, green squares: 0.6 mM, and blue triangles: 1.0 mM). Dashed lines are fitting curves using *p*_*u*_ = *p*_*uo*_ − *k* ⋅ *c*, where *p*_*u*_ is the percentage of unbound DNA and *c* is the concentration of H-NS proteins. (d) Dependence of the fitted slopes (*k*) from panel c on the concentration of Ag^+^ ions. Error bars represent fitting errors.

We quantified the intensities of the unbound DNA bands, from which the percentages of unbound DNA were calculated, 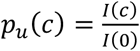, where *I*(*c*) is the intensity of unbound DNA band in the presence of H-NS proteins at a concentration of *c* and *I*(0) is the intensity of unbound DNA band without H-NS proteins on the same gel. At constant concentrations of Ag^+^ ions (i.e., comparing bands in the same gel in Fig. 2a), the percentage of unbound DNA decreased linearly as the concentration of H-NS proteins increased (Fig. 2c), confirming the binding of H-NS proteins on DNA. At constant concentrations of H-NS proteins (i.e., comparing bands in the last lanes of different gels in Fig. 2a), the percentage of unbound DNA increased steadily as the concentration of Ag^+^ went up from 0 to 1 mM (Fig. 2b). In other words, Ag^+^-treatment led to less DNA to be bound by the H-NS proteins.

It is worthwhile to point out a key difference between our EMSA assay and conventional EMSA assays commonly used in the literature ^[45–47]^: the concentrations of DNA and proteins were not around the dissociation constant (*K*_*D*_). The rationale for our non-conventional EMSA assay is three-fold. First, the conventional EMSA assay is not suitable for our purpose in this study. Instead of measuring the absolute binding affinity between H-NS proteins and DNA (usually reported by the dissociation constant), it was desired to compare the effect of Ag^+^ ions on the binding affinity. Therefore, the experiments were designed so that all the conditions (i.e., the concentrations of DNA and proteins), except the concentration of Ag^+^ ions, were kept constant. Second, the kinetics from the binding reaction equation is valid and can be analytically solved for the concentrations of DNA and proteins that are far from *K*_*D*_. For the binding of H-NS proteins (P) on DNA (D), D + P ⇌ DP, the concentration of unbound DNA is simply

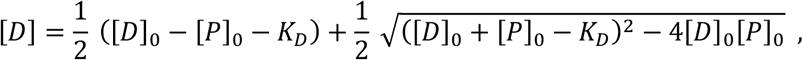

where [*D*]_0_ and [*P*]_0_ are the initial concentrations of DNA and proteins, respectively. Third, the binding reaction equation predicts that the dependence of the amount of unbound DNA [*D*] on the initial concentration of proteins [*P*]_0_ (low enough) is roughly linear (in linear scale) for a wide range of dissociation constants *K*_*D*_, [*D*] ≈ [*D*]_0_ − *k*[*p*]_0_ (Fig. S5a). Although an analytical relation between *k* and *K*_*D*_ is not trivial, this parameter [is negatively related to the dissociation constant *K*_*D*_ (the higher *K*_*D*_ is, the lower *k* is, as shown Fig. S5b). As a result, the negative slopes *k* could be used to equivalently report the binding affinity of H-NS proteins on DNA. The negative slopes *k* were extracted by fitting the EMSA data with lines (Fig. 2c). We observed that *k* decreased as the concentration of Ag^+^ ions increased (Fig. 2d). This observation suggested that the dissociation constant *K*_*D*_ increased at higher concentration of Ag^+^ ions, again indicating that Ag^+^ ions weakened the binding of H-NS proteins on duplex DNA.

### Dehybridization of bent duplex DNA induced by Ag^+^ ions

Our results showed that Ag^+^ ions interact weaken the binding of H-NS proteins on double-stranded DNA. A further question is how Ag^+^ ions affect the binding between DNA and H-NS proteins. It is possible that both DNA and H-NS interact with Ag^+^ ions, as suggested by the previous studies. First, DNA has also been reported to interact with Ag^+^ ions. In addition to the electrostatic interactions, Ag^+^ ions bind to cytosine–cytosine (C–C) mismatching base pair selectively ^[48–50]^ and possibly result in chain-slippage ^[51]^. Second, it has been reported that Ag^+^ ions interact with thiol groups in proteins (e.g., cysteine) and peptides containing motifs of HX_n_M or MX_n_H (H – histidine, M – methionine, X – other amino acids) ^[52–54]^. On the other hand, the H-NS protein does not contain any HX_n_M or MX_n_H motifs but has a single cysteine in the dimerization domain, instead of the linker and DNA-binding domain ^[36,55]^; therefore, the interaction of H-NS and Ag^+^ ions is expected to minimally affect the binding of the protein on DNA. Based on these previous results, we hypothesize that Ag^+^ ions affect the DNA hybridization, resulting in partial dehybridization (i.e., dissociation of double-stranded DNA into single strands), or tendency towards dehybridization, and weakening the binding between DNA and H-NS proteins.

Direct interaction between DNA and Ag^+^ ions was confirmed by isothermal titration calorimetry (ITC) using short linear double-stranded DNA of 25 base pairs. Isothermogram representing the Ag^+^-DNA binding suggest that the metal ions interact with DNA *albeit* weakly (Fig. 3A). The Ag^+^-DNA is exothermic and proceeds with modest evolution of heat. The heat exchanges of first few injections were observed more than four times of that of the controls (Figs. 3B – 3D) and then decrease to reach plateau at ~25 injections (Fig. S6). Although the two strands are fully complementary to each other, four possible C-C mis-pairs may be formed by one-base-slippage, mediated by Ag^+^ ions. Therefore, this result seems to corroborate with the number of possible mis-pair C-C in the DNA ^[48–50]^. Fitting the ITC data using the one set of sites binding model (i.e., assuming all the binding sites on the DNA to Ag^+^ ions are equal and have the same binding affinity) gave a binding constant of *K* = (5.29 ± 3.37) × 10^4^ M^−1^, indicating a weak interaction between the Ag^+^ ions and the DNA, and a heat change of Δ*H* = −12.63 ± 3.66 kcal/mol, confirming that the binding of Ag^+^ ions on the DNA was an exothermic process (Fig. S6). It is noted that the non-specific electrostatic interactions between the negatively charged DNA and the Ag^+^ cations ^[56]^ are expected to contribute to the ITC results, which complicate the further quantitative analysis of DNA-ion interactions ^[57]^.

**Figure 3.**
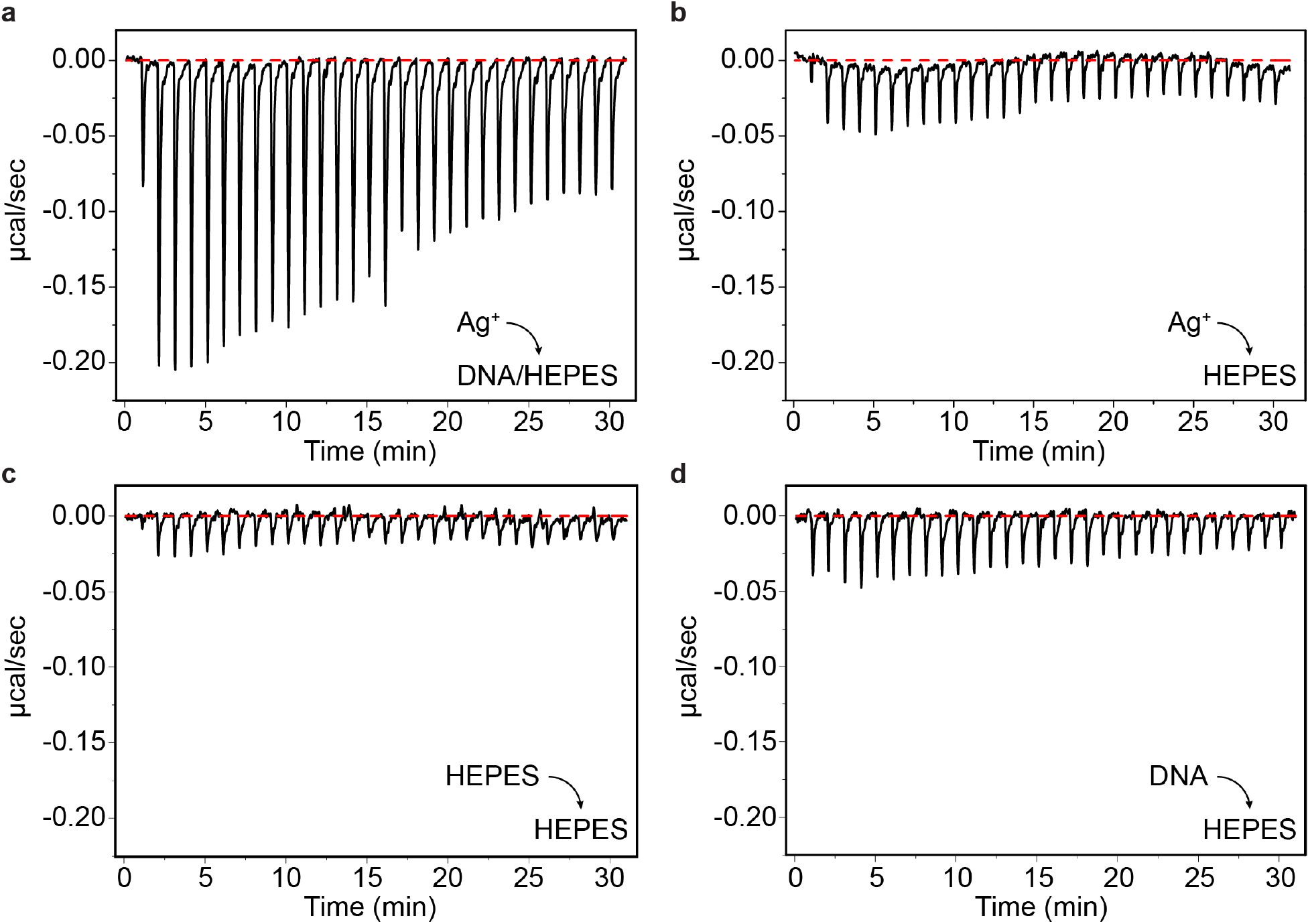
Direct interaction between Ag^+^ ions and double-stranded DNA measured by isothermal titration calorimetry (ITC), in which 0.2 mM Ag^+^ ions were titrated into DNA at pH 7.4 in 0.2 mM tris-HCl and 0.25 mM NaCl at 25 °C in 10 mM HEPES buffer: (a) 1 µM DNA; and (b) no DNA. The injection volume was 1.3 µL with 1 min interval between injections. Additional control experiments were performed by titrating (c) HEPES buffer or (d) 1 µM DNA into HEPES buffer.

Next, we attempted to directly probe the dissociation of double-stranded DNA into single strands induced by Ag^+^ ions, which remains a challenge for two reasons. First, direct interactions between Ag^+^ ions and DNA might not be strong enough to open up the double strands under normal conditions. Second, there exist competing effects of Ag^+^ ions, including electrostatic interactions between positively charged Ag^+^ ions and negatively charged DNA, which are expected to stabilize the double-stranded DNA. As a result, Gogoi et al. ran gel electrophoresis on plasmid DNA from Ag-treated bacteria but no direct effects were observed ^[58]^. In addition, when we treated short, linear, double-stranded DNA with Ag^+^ ions at concentrations ranging from 0 to 90 µM, we did not observed any dehybridization of the linear double-stranded DNA (Fig. 4b) and the intensities for the bands of the double-stranded DNA did not change significantly (red squares in Fig. 4d) ^[59]^.

**Figure 4.**
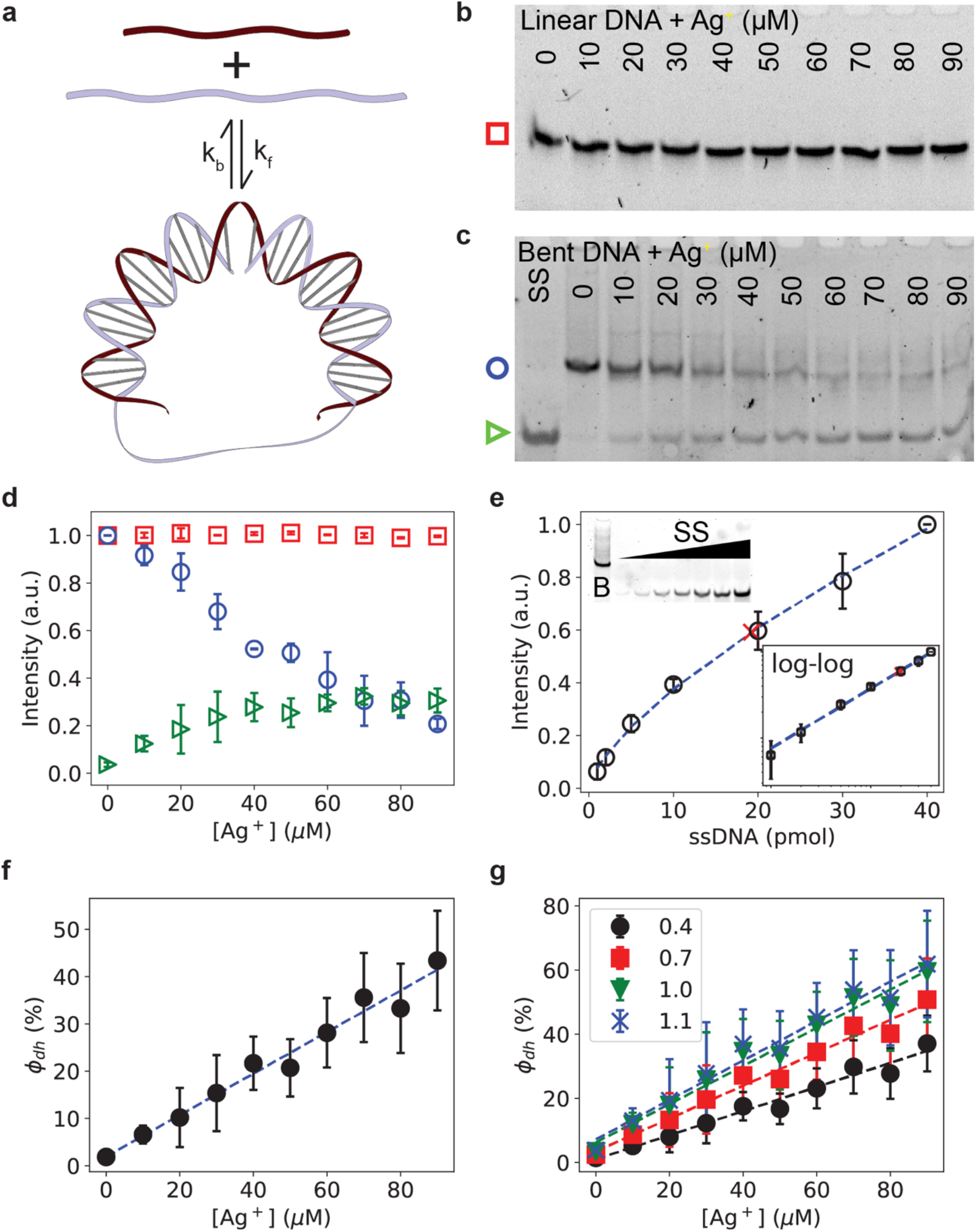
Dehybridization of bent double-stranded DNA induced by Ag^+^ ions. (a) Self-assembly of a circular bent double-stranded DNA that was used in this study to amplify and probe the effect of Ag^+^ ions on double-stranded DNA. (b) An example gel of linear double-stranded DNA in the presence of 0-90 µM Ag^+^ ions. (c) An example gel of bent double-stranded DNA in the presence of 0-90 µM Ag^+^ ions. (d) Dependence of normalized intensities of the bands from panels b and c on the concentration of Ag^+^ ions (0-90 µM). The symbols in this plot correspond to bands indicated by the same symbols in panels b and c. (e) Fluorescence intensity of bands of single-stranded DNA (*I*_*SS*_) stained by Sybr Safe as a function of their amount (*x*_*SS*_, black circles), which was fitted by 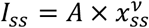 (blue dashed curve, *v*=0.69±0.02). Based on the intensity of the band of the bent double-stranded DNA at 10 pmol (red cross) on the same gel, the “equivalent” amount of the bent DNA was determined from the fitted equation and used to estimate the correction factor, *β* ≈ 0.52. Left-top inset: bent double-stranded DNA of 10 pmol and single-stranded DNA of varying amounts (1-40 pmol) on the same gel. Right-bottom inset: the same data in log-log scale. (f) Dependence of the fraction of dehybridization (*φ*_*dh*_) on the concentration of Ag+ ions (0-90 µM) for bent double-stranded DNA, 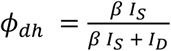 where *β*=0.52 is the measured correction factor, and *I*_*s*_ and *I*_*D*_ are the normalized intensities of the bands for the single-stranded DNA due to dehybridization (indicated by the green triangle in panel c) and for the bent double-stranded DNA (indicated by the blue circle in panel c), respectively. Linear fitting of the data (blue dashed line) resulted in a slope of (0.44±0.05)% per µM. (g) Dependence of the fraction of dehybridization (*φ*_*dh*_) on the concentration of Ag^+^ ions (from 0 to 90 µM) for bent double-stranded DNA using different correction factors: *β*=0.4 (black circles), 0.7 (red squares), 1.0 (green triangles) and 1.1 (blue crosses). The dashed lines are linear fitting, resulting in slopes ranging from (0.38±0.02)% per µM (with *β*=0.4) to (0.61±0.04)% per µM (with *β*=1.1).

To overcome this challenge to test our hypothesis, we exploited our recently developed method using bent DNA molecules ^[59]^. In this method, two single strands of DNA of different lengths (45 bases and 30 bases) form a circular bent DNA molecule upon hybridization ^[60–66]^ (Fig. 4a). Stresses in the circular DNA due to the bending of the double-stranded segment make the molecule more prone to perturbations and thus “amplifier” interactions between the DNA with other molecules ^[59]^. The rationale of using the bent DNA molecules is three-fold. First, this new method can amplify weak interactions between DNA and other molecules, and thus it makes it easier to detect the possible interactions ^[59]^. Second, the possible DNA dehybridization induced by Ag^+^ ions may be directly and conveniently visualized by gel electrophoresis. Third, it has been reported that H-NS proteins bind to curved DNA and H-NS proteins facilitate bridging of DNA strands ^[46,67,68]^; therefore, bent DNA might mimic the natural DNA that H-NS proteins bind in live bacteria more effectively than linear DNA.

We observed that Ag^+^ ions caused the intensity of the bent DNA band to decrease (Fig. 4c). Additionally, bands ahead of the bent DNA band showed up in the presence of Ag+ ions (indicated by the green triangle in Fig. 4c). On the same gel, we included a lane for the longer single-stranded DNA (45 bases) and found that the bands appeared in the presence of Ag^+^ ions matched with this single-stranded DNA band very well, suggesting that Ag^+^ ions led to dehybridization of the circular bent DNA. It is worthwhile to note that the dehybridization of the bent DNA was observed at a concentration of Ag^+^ ions as low as 10 µM (Fig. 4c). The observations were quantified by measuring the intensities of the bands in Fig. 4c, which were normalized to the intensity of the bent DNA band in the absence of Ag^+^ ions. The intensities of the bands for the bent double-stranded DNA (*I*_*D*_) decreased steadily as the concentration of Ag^+^ ions increased, while the intensities of the bands for the single-stranded DNA (*I*_*S*_) increased after Ag^+^-treatment. We further estimated the percentage of dehybridization of the bent DNA caused by Ag^+^ ions using 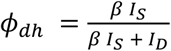, where *β* is a correction factor to account for differences in the staining efficiencies of Sybr Safe dyes for double-stranded DNA and single-stranded DNA, respectively. This factor was measured in a previous study and ranged from 0.4 to 1.1 ^[69]^. Following Ref. ^[69]^, we experimentally measured this correction factor by staining both bent double-stranded DNA and single-stranded DNA on the same gel (left-top inset of Fig. 4e). We varied the amount of single-stranded DNA (SS bands) but kept the amount of bent double-stranded DNA (B band) constant. From the fluorescence intensity (*I*_*SS*_) of bands of single-stranded DNA stained by Sybr Safe as a function of their amount (*x*_*SS*_), we obtained a “calibration” curve (Fig. 4e). Interestingly, the fluorescence intensity was not linear to the amount of single-stranded DNA, presumably due to the background intensities of the gel. Instead, the “calibration” curve could be fitted well with a power-law, 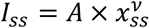, which could be seen more clearly from the log-log plot (Fig. 4e). The fitting resulted in m = 0.69 ± 0.02 (Fig. 4e). From the fitting, we estimated the “equivalent” amount of the bent double-stranded DNA at 10 pmol from its intensity (Fig. 4e). By comparing the “equivalent” amount and the actual amount of the bent double-stranded DNA, we determined that the correction factor in our experiments was *β* ≈ 0.52, which fell in the previously reported range of [0.4, 1.1] ^[69]^. Using the measured correction factor, we estimated the percentage of dehybridization *φ*_*dh*_ at different concentrations of Ag^+^ ions and observed that *φ*_*dh*_ was roughly linear to the concentration of Ag^+^ ions, with a slope of [(0.44±0.03)% per µM] (Fig. 4f). We also note that the linear dependence was robust. For example, when we varied the correction factor from 0.4 to 1.1, the *φ*_*dh*_-vs-[Ag^+^] plots remained linear for different correction factors (Fig. 4g). On the other hand, the slopes varied from [(0.38±0.02)% per µM] for *β*=0.4 to [(0.61±0.04)% per µM] for *β*=1.1.

### A plausible mechanism for the faster dynamics of H-NS proteins caused by Ag^+^ ions

Based on our experimental results and analyses using various biological assays, we proposed the following mechanism for the effects of Ag^+^ ions on the diffusive dynamics of H-NS proteins in live bacteria. The Ag^+^ ion interaction with DNA leads to a tendency of dehybridization (or a partial dehybridization) of double-stranded DNA (Figs. 5a and 5b). The partial dehybridization is likely amplified in the segment of the curved DNA where H-NS proteins preferably bind, and therefore weakens the binding of H-NS proteins on the bacterial genome. The weakened binding results in an increasing fraction of unbound H-NS in the bacteria (Figs. 5b and 5c). As unbound H-NS proteins diffuse faster than the bound ones, the overall diffusive dynamics of H-NS proteins became faster after Ag^+^-treatment.

**Figure 5.**
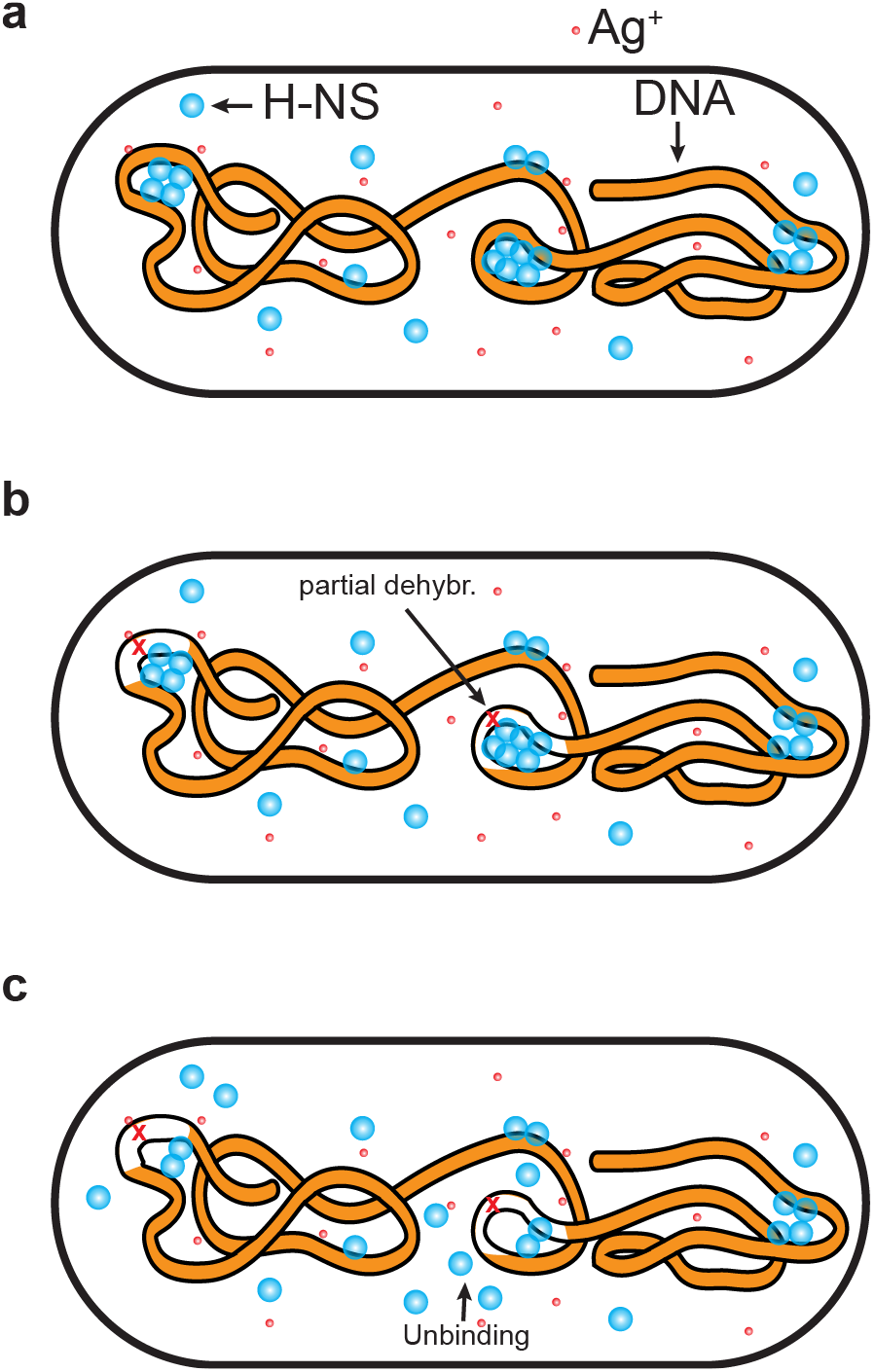
Speculated mechanism of Ag^+^ ions’ effects on the diffusive dynamics of H-NS proteins in live bacteria. (a) A bacterium subjecting to Ag^+^ ions. (b) Partial dehybridization of curved DNA induced by Ag^+^ ions. (c) Unbinding of H-NS proteins from DNA due to (partial) dehybridization, leading to faster diffusive dynamics of the H-NS proteins.

## Discussion

We used super-resolution fluorescence microscopy in combination with single-particle tracking to investigate the diffusive dynamics of H-NS proteins in live bacteria treated with Ag^+^ ions. We observed that Ag^+^-treatment led to faster dynamics of H-NS proteins: while the motion of H-NS proteins remains sub-diffusive, the generalized diffusion coefficient of H-NS proteins increased upon exposure to Ag^+^ ions. To understand the mechanism of the observed faster dynamics of H-NS, electrophoretic mobility shift assay was performed *in vitro* with purified H-NS proteins and double-stranded DNA. It was found that Ag^+^ ions weaken the binding between H-NS proteins. With isothermal titration calorimetry, we confirmed that DNA and Ag^+^ ions interact directly. Furthermore, we examined the effect of the DNA-Ag interaction using our recently developed sensors based on bent DNA molecules, and found that Ag^+^ ions caused dissociation of double-stranded DNA into single strands. Our results suggest a plausible mechanism for the faster dynamics of H-NS proteins in live bacteria when subjected to Ag^+^ ions due to Ag^+^-induced DNA dehybridization.

The observed faster diffusion of H-NS proteins in bacteria upon exposure to Ag^+^ ions was unexpected because the metal ion, due to its antimicrobial effects, is likely to reduce the metabolic rate of bacteria, lower the fluidity of bacterial cytoplasm, and slow down the diffusion of proteins in bacterial cytoplasm ^[70]^. Similar effects with other antibiotics have been observed previously ^[29,70,71]^. However, due to the specific functions of H-NS proteins (e.g., binding to DNA) ^[34]^, the change in the diffusion of H-NS proteins was opposite to expectations upon exposure to Ag^+^ ions. This unexpected observation raises awareness in the field of understanding of the material and physical properties of biological systems at the cellular level. As many current studies on the mechanical properties of bacterial and cellular cytoplasm are based on monitoring the motion and diffusion of tracers (proteins or other molecules/particles) in the organisms of interest ^[70–81]^, it is important to pay close attention to the function of these tracer molecules/particles. As evidenced in the literature, molecules with different functions display different diffusive behaviors in *E. coli* bacteria ^[32,70–79]^, which would be translated to the differences in the material properties of the bacterial cytoplasm experienced by the molecules. In addition, this function-dependence indicates another contributing factor to the heterogeneity of the physical properties of cellular cytoplasm.

To our knowledge, this work presents the first study of the antimicrobial effects of Ag^+^ ions on the diffusive dynamics of proteins at the molecular level in live bacteria. H-NS is one major member of the ≥12 nucleoid-associated proteins (NAPs) in gram-negative bacteria ^[34,82]^. In addition, many fundamental cellular processes in bacteria and cells rely on interactions between DNA and proteins, including DNA packaging ^[83]^, gene regulation ^[34,82,84,85]^ and DNA repairing ^[86–88]^. It remains unclear how the diffusive dynamics of these DNA-interacting proteins are affected by Ag, and whether the effects of Ag^+^ ions on the other DNA-interacting proteins are different from that of H-NS proteins.

Dissociation of DNA and DNA-binding proteins has long been reported in the literature. For example, DNA were dissociated from the histone proteins and released from the nucleosomes in the presence of salt solutions (i.e., ions) at high concentrations (e.g., ~750 mM NaCl), which was attributed to the electrostatic screening effect of the ions on the negative charges of the DNA backbone ^[89]^. However, it is important to point out that the electrostatic effect is unlikely the major contributor to the dissociation of DNA and H-NS proteins observed here because of the low concentration of Ag^+^ ions (10 µM) used in the current study. An interesting future study would be to understand how the nucleosome core particles are affected by Ag^+^ ions, which is expected to shed light on possible mechanisms of cytotoxicity of Ag on eukaryotic cells.

It is worthwhile to emphasize that the current study does not exclude the possibility that interactions between H-NS proteins and Ag^+^ ions affect the binding between H-NS and DNA. For example, although the only cysteine in H-NS for the Ag-thiol interactions is in the dimerization domain, the binding affinity of this protein on DNA could be changed allosterically. Allosteric regulation (or allosteric control) – the regulation of a protein by molecules at a site other than the protein’s active site ^[90–92]^ – was well-known in regulatory proteins such as lactose repressor ^[90,91]^. It would be interesting to further investigate whether and how the binding of H-NS proteins on DNA is allosterically regulated by their interactions with Ag^+^ ions.

Previous studies have suggested that Ag^+^ ions cause serious damages in the cell membrane in various aspects, such as detachment of inner membrane from the outer envelope and cis/trans transformation of the unsaturated membrane fatty acids ^[17,93–97]^. However, a dynamic picture of the Ag-caused membrane damages is still missing. Interesting questions include how the fluidity of membrane lipids is affected and how the membranes are disrupted by Ag. Dynamic studies with both high spatial and temporal resolutions are required to address these questions. It would be exciting to apply the methodology described in this work to answer these questions.

## Methods

### Bacterial strain and sample preparation

The *E. coli* strain used in this study is JW1225 of the Keio collection ^[98]^ (purchased from the Yale *E. coli* Genetic Stock Center) transformed with a plasmid pHNS-mEos3.2, which encodes *hns-meos* fusion gene ^[32,99]^. The resultant strain expresses H-NS proteins fused to mEos3.2 photo-switchable fluorescent proteins ^[99,100]^ and carries resistance against kanamycin and chloramphenicol ^[32,99]^. The same strain has been used in our previous studies ^[32,43]^.

The bacteria were grown overnight in a defined M9 minimal medium, supplemented with 1% glucose, 0.1% casamino acids, 0.01% thiamine and appropriate antibiotics (kanamycin + chloramphenicol) at 37ºC in a shaking incubator with a speed of 250 rpm ^[32,43,101,102]^. On the next day, the overnight culture was diluted 50 to 100 times into a fresh medium such that OD_600_ = 0.05. This culture (5 mL) was regrown in the shaking incubator at 37ºC for 2-3 hr. When the OD_600_ of the bacterial culture reached 0.3, 10 µL of the culture were transferred onto a small square of agarose gel pad (5 mm x 5 mm). The remaining bacterial culture was treated with prepared stock solution of Ag^+^ ions (final concentration = 10 µM). The stock solutions of Ag^+^ ions were prepared by dissolving AgNO_3_ powders (Alfa Aesar, Haverhill, MA) in deionized water (>17.5 MΩ), followed by filtration and stored at 4ºC in dark for later use. The Ag^+^-treated bacteria were incubated at 37ºC in the shaking incubator (250 rpm) for 2, 4, 6, and 8 hr. After each two hours, 10 µL of the bacterial culture were taken from the treated culture and added into new square of agarose pads containing Ag^+^ ions at 10 µM. The control (untreated) and treated samples were left in dark at room temperature for 20-30 minutes on the agarose pad to allow the bacteria to be absorbed and mounted. The agarose pad was then flipped and attached firmly and gently to a clean coverslip, which was glued to a rubber O-ring and a clean microscope slide to form a chamber ^[32,43]^.

### Single-particle tracking photoactivated localization microscopy (sptPALM) on H-NS proteins in live bacteria

The super-resolution fluorescence microscope used in this work is home-built based on an Olympus IX-73 inverted microscope with an Olympus oil immersion TIRF objective (100X N.A. = 1.49). The microscope and data acquisition were controlled by Micro-Manager ^[103]^. To activate and excite the H-NS-mEos3.2 fusion proteins in live *E. coli* bacteria, lasers at 405 nm and 532 nm from a multilaser system (iChrome MLE, TOPTICA Photonics, NY, USA) were used ^[32,43,100]^. Emissions from the fluorescent proteins were collected by the objective and imaged on an EMCCD camera (Andor, MA, USA) with an exposure time of 30 ms, which resulted in 45 ms for the actual time interval between frames. The effective pixel size of acquired images was 160 nm. For each sample (untreated or treated for 2-8 hr), 5-8 movies were acquired.

The resulting movies (20,000 frames) were analyzed with RapidStorm ^[104]^, generating x/y positions, x/y widths, intensity, and background for each detected fluorescent spot. Spots with localization precision >40 nm were rejected ^[25,32]^. The spots that survived the criteria were further corrected for drift using a mean cross-correlation algorithm ^[105]^. Furthermore, the spots were segmented manually into individual cells. The positions ***r*** from the same molecule in adjacent frames in the same cells were linked by standard algorithms with a memory of one frame (Mem=1) and a maximum step size of 480 nm (Maxdisp=480) using trackpy ^[27,29,106,107]^, from which the trajectories of individual molecules ***r***(*t*) were obtained. Velocities of H-NS proteins were then calculated from the trajectories, 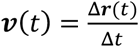, where Δ***r***, is the displacement and Δ*t* is the time interval between adjacent data points in the trajectories.

### Ensemble mean-square-displacement (eMSD) and generalized diffusion coefficient

From the trajectories, **r**(*t*) in each bacterial cell, the ensemble mean-square-displacements (eMSD) were calculated 〈Δ*r*^2^(*τ*)〉 = 〈(***r***(*t* + *τ*) − ***r***(*t*))^2^〉 using built-in functions in trackpy ^[107]^. The eMSD data were then averaged over different cells from multiple movies for the same sample. The number of bacterial cells ranged from 158 to 678. The averaged eMSD data were then fitted using 〈Δ*r*^2^(*τ*)〉 = 4*Dτ*^*α*^, resulting the generalized diffusion coefficient (*D*) and the anomalous scaling exponent α.

### Plasmid cloning for H-NS expression and purification

A plasmid – pHisHNS – was constructed for the expression and purification of H-NS proteins for *in vitro* experiments. Briefly, the *hns* gene was amplified from the pHNS-mEos3.2 plasmid by polymerase chain reaction (PCR) using a pair of primers (H-NS-F: 5’-GGG GAC AAG TTT GTA CAA AAA AGC AGG CTC CAT GAG CGA AGC ACT TAA-3’ and H-NS-R: 5’-GGG GAC CAC TTT GTA CGG GAA AGC TGG GTT TTA TTG CTT GAT CAG GAA-3’). The PCR fragment was cloned into the entry vector pENTR/D-TOPO (Thermo Fisher Scientific, MA, USA) using BP Clonase enzyme mix (Thermo Fisher Scientific, MA, USA), resulting in an entry clone. The entry clone was mixed with the pDEST^TM^17 vector and LR Clonase enzyme mix (Thermo Fisher Scientific, MA, USA), generating the plasmid pHisHNS, which encodes 6xhis-tagged H-NS proteins. The final plasmid was verified by PCR and sequencing (Eton Bioscience Inc., CA, USA).

### Expression and purification of H-NS proteins

The constructed plasmid, pHisHNS, was used for expression of H-NS proteins. Briefly, it was transformed to *E. coli* BL21(DE3) competent cells (New England Biolabs, MA, USA). On the second day, a single colony was inoculated into 15 mL of LB medium and grown at 37ºC in a shaking incubator (250 rpm) for overnight. The overnight culture was transferred into 400 mL fresh ^LB medium, and regrown at 37ºC to reach OD_600_ = 0.4, followed by induction with 0.5mM isopropyl^ β-D-1-thiogalactopyranoside (IPTG) (IBI Scientific, IA, USA) at 30ºC for 5 hr. Cells were collected by centrifuging at 4500 rpm at 4ºC for 30 min, and the cell pellets were stored at −80ºC.

On the next day, the cell pellets were resuspended in 30 mL of lysis buffer (1X PBS with 10 mM imidazole) containing 1mM phenlymethanesulfonyl fluoride (PMSF) (Bio Basic, Canada) and 1X protease inhibitors (Roche, Switzerland). The resuspended cells were then further lysed by sonication on ice (30% pulse for 10 sec and rest for 15 sec, repeated for 3 min), followed by mixing with 0.1% Triton X-100 and incubation with shaking at 4ºC for 1 hr. The cell lysate was centrifuged at 12,000 rpm for 30 min, from which the supernatant was collected and filtered with syringe (0.45 µm). The filtered supernatant was mixed with nickel-beads at 50% bed volume (Thermo Fisher Scientific, MA, USA), and incubated at 4ºC for overnight. On the next day, the mixture was applied to a Poly-Prep Chromatography Columnscolumn (Bio-Rad Laboratories, CA, USA) and washed with wash buffer I (lysis buffer with 10 mM imidazole) twice and wash buffer II (lysis buffer with 20 mM imidazole) twice. After the extensive washes, the his-tagged H-NS proteins were eluted from the nickel beads by 5 mL elution buffer (lysis buffer with 250 mM imidazole). The eluted proteins were concentrated using Amicon concentrators with a cut-off of 10 kDa (Millipore Sigma, MA, USA). The purity of the purified his-tagged H-NS proteins was measured using SDS-PAGE (15%) gel electrophoresis. The concentration of the purified protein was measured by Bradford assay ^[108,109]^.

### Electrophoretic mobility shift assay for binding of H-NS proteins on DNA

Electrophoretic mobility shift assay (EMSA) ^[45]^ was used to examine the binding of H-NS proteins on DNA in the absence and presence of Ag^+^ ions. Briefly, 1 uL of double strand DNA (~300 bp at ~0.375 µg/µL) was mixed with purified H-NS proteins of various volumes in binding buffer (10 mM Tris at pH 7.5, 15% glycerol, 0.1 mM EDTA, 50 mM NaCl, and 1 mM 2-mercaptoethanol) with a total volume of 20 µL. The volume of H-NS proteins (stock concentration: 0.65 µg/µL) ranged from 0 to 10 µL, resulting in a final concentration of the H-NS protein ranging from 0 to 433 µM. The mixtures of proteins and DNA were incubated on ice for 15 min and then at room temperature for 30 min. To examine the effect of Ag^+^ ions on the binding of H-NS proteins on DNA, the samples were prepared the same as the negative control, except that the binding buffer contained Ag^+^ ions at final concentrations of 0.1, 0.6, or 1 mM. Following the incubation, the samples were mixed with DNA loading buffer (Bio-Rad Laboratories, CA, USA) and subjected to PAGE (3%) gel electrophoresis in 1X TBE buffer at 100 V for 50 min. The gels were stained with Sybr Safe (Thermo Fisher Scientific, MA, USA), and imaged by a ChemiStudio gel documentation system (Analytik Jena, Germany). The gel images from the EMSA assays were analyzed using ImageJ ^[110,111]^. Each set of samples were repeated for at least three times, from which the averages and the standard errors of the means (SEM) were calculated.

### Isothermal titration calorimetry (ITC) measurements

The ITC measurements were carried out at 25 °C using an isothermal titration calorimeter (MicroCal iTC200, Malvern) equipped with a 280 µL sample cell and a pipette syringe with a spin needle. The sequences of the double-stranded DNA are 5’-GTG CTG ACG GAA TTC TTG ACA TCT C-3’ and 5’-GAG ATG TCA AGA ATT CCG TCA GCA C-3’. For the experiment, 250 µL of 1 µM DNA in 10 mM HEPES buffer (pH 7.5) containing 0.2 mM Tris-HCl and 0.25 mM NaCl was placed in the cell. Then, 0.2 mM AgNO_3_ in 10 mM HEPES buffer was titrated to the cell using 1.3 µL per injection and 1 min interval between injections. During the titration, a spin rate of 750 rpm was used to mix the reactants. For the control experiment, 10 mM HEPES buffer containing 0.2 mM Tris-HCl and 0.25 mM NaCl in the absence of DNA was placed in the cell instead for titration.

### Probing the DNA-Ag^+^ interactions using bent DNA amplifiers

Bent DNA amplifiers were prepared as described previously ^[59,61,63,64]^. Briefly, synthesized single-stranded DNA molecules (Integrated DNA Technologies, IL, USA) were resuspended in distilled water to a final concentration of 100 µM. The sequences of DNA strands for constructing bent DNA molecules are 5’-CAC AGA ATT CAG CAG CAG GCA ATG ACA GTA GAC ATA CGA CGA CTC-3’ (long strand, 45 bases) and 5’-CTG CTG AAT TCT GTG GAG TCG TCG TAT GTC-3’ (short strand, 30 bases). Single strands were mixed at equal molar amount in background buffer (0.4 mM Tris-HCl and 0.5 mM NaCl, pH=7.5) containing Ag^+^ ions at various concentrations ([Ag^+^] = 0, 10, 20, …, 80, 90 µM). Ag^+^ ions were provided from aqueous solutions of AgNO_3_. The final concentrations of the DNA were 2 µM. The mixtures were heated to 75ºC for 2 minutes, and gradually cooled down to 22ºC (room temperature) in 5 hours. Upon hybridization, a circular construct is formed, with a double-stranded portion of 30 bp (with a nick) and a single-stranded portion of 15 bases (Fig. 4a). The mixtures were incubated at 22ºC for overnight to allow full equilibrium, followed by PAGE gel electrophoresis. Experiments were performed in triplicates. Imaging and analysis of the DNA gels were performed similarly as in the EMSA assay described above.

## Acknowledgment

We thank Dr. Joshua N. Milstein for the generous gift of the pHNS-mEos3.2 plasmid. This work was supported by the University of Arkansas, the Arkansas Biosciences Institute (Grant No. ABI-0189, ABI-0226, ABI-0277, ABI-0326), and the National Science Foundation (Grant No. 1826642). We are also grateful for supports from the Arkansas High Performance Computing Center (AHPCC), which is funded in part by the National Science Foundation (Grants No. 0722625, 0959124, 0963249, 0918970) and the Arkansas Science and Technology Authority.

## Author Contributions

YW designed the project and experiments. AAS and VK performed sptPALM experiments. PK cloned plasmids for expression and purification of H-NS proteins. PK and AAS performed EMSA experiments. RKG, RM, PK, JF and MR performed ITC experiments. JF performed experiments using bent DNA. All the authors performed data analysis. YW, AAS, PK and JC wrote the manuscript. All the authors edited and reviewed the manuscript.

## Additional Information

**Supplementary Information** is available for this paper at https://doi.org/10.xx/xx.xx/xx.xx

