## Supplementary Information for "Faster diffusive dynamics of histone-like nucleoid structuring proteins in live bacteria caused by silver ions"

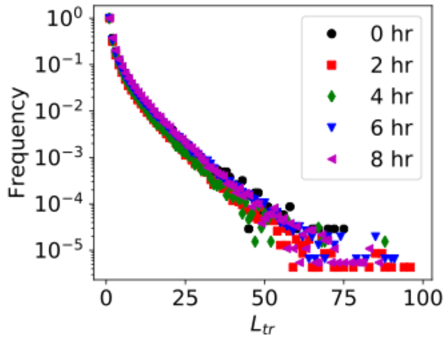


**Figure S1**. Distributions of the lengths of trajectories of H-NS proteins in bacteria before $Ag^+$-treatment (0 hr) and after $Ag^+$-treatment for 2 to 8 hr.

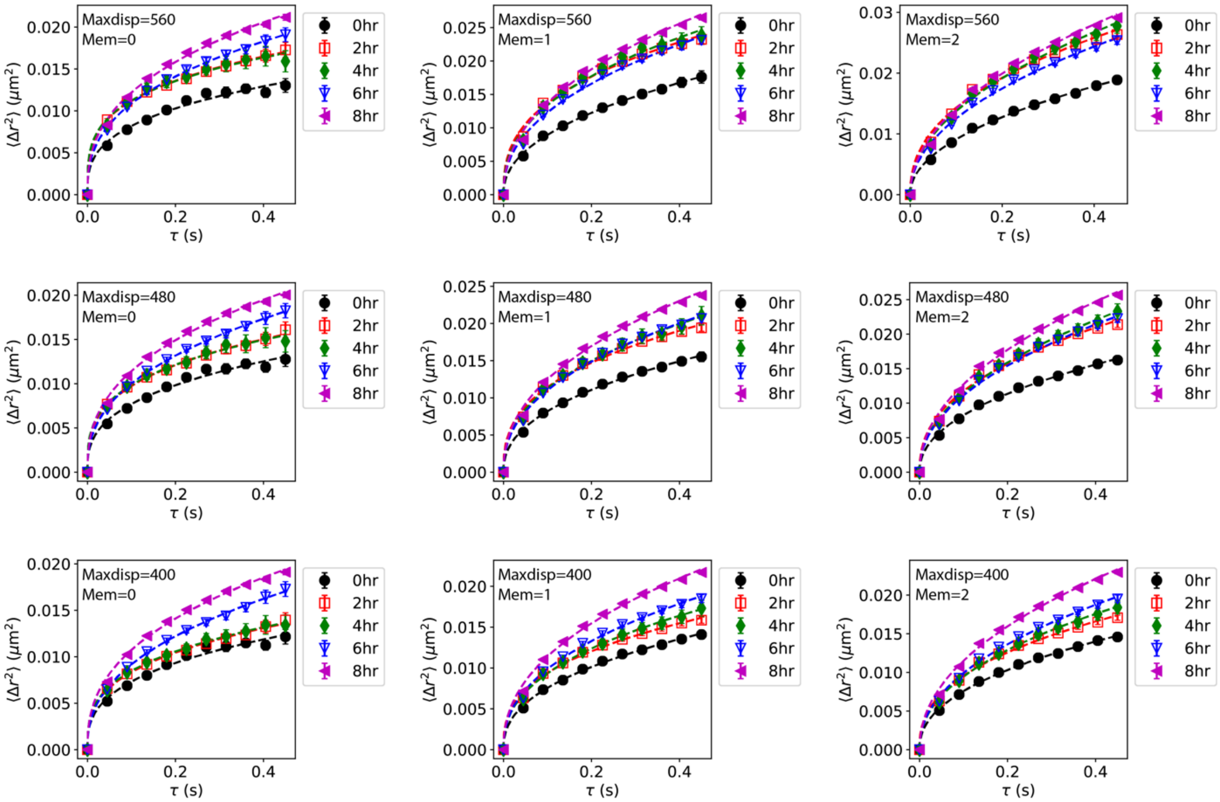


**Figure S2**. Effects of varying tracking parameters, Maxdisp and Mem, on the eMSD curves.

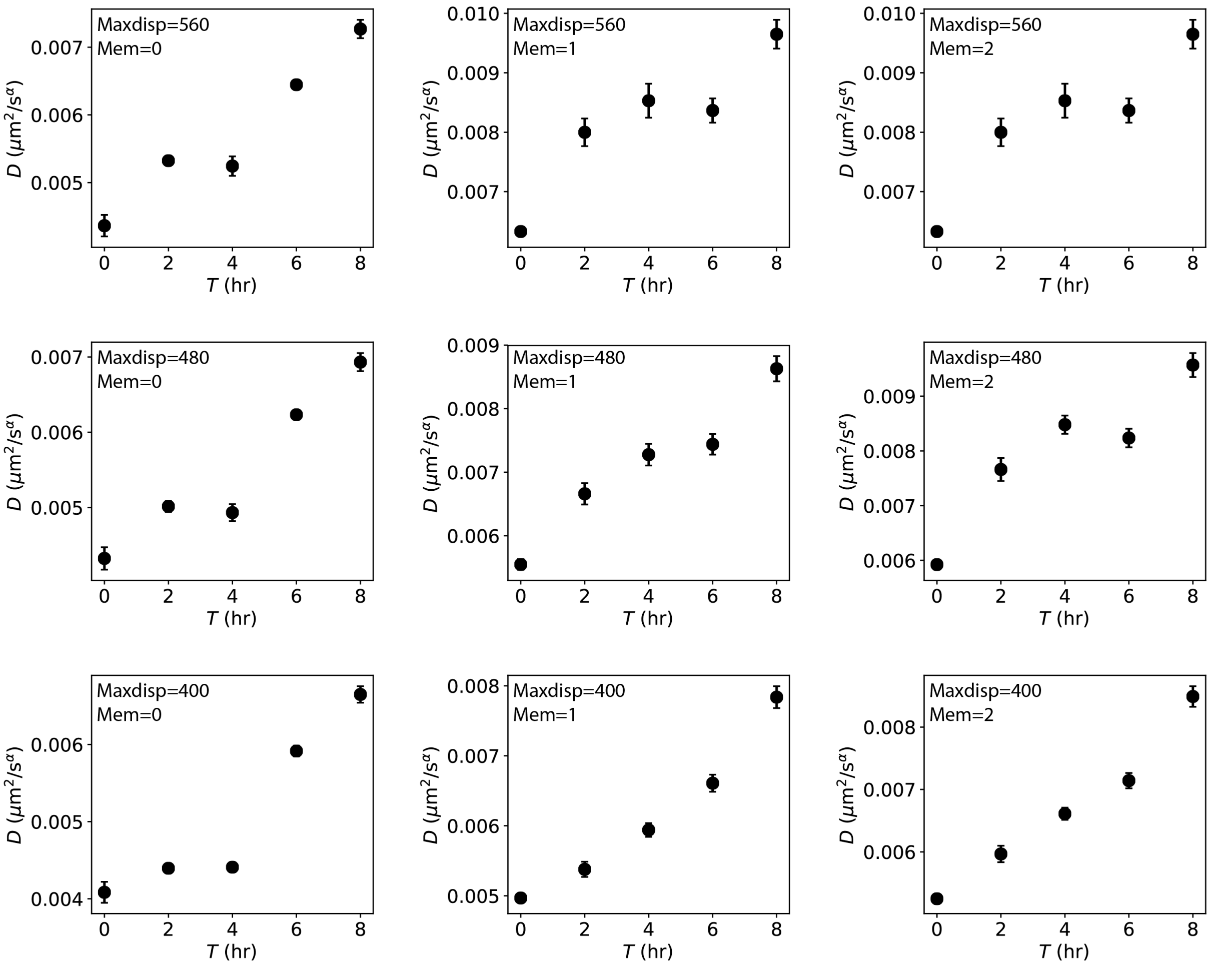


**Figure S3**. Effects of varying tracking parameters, Maxdisp and Mem, on the fitted generalized diffusion coefficients.

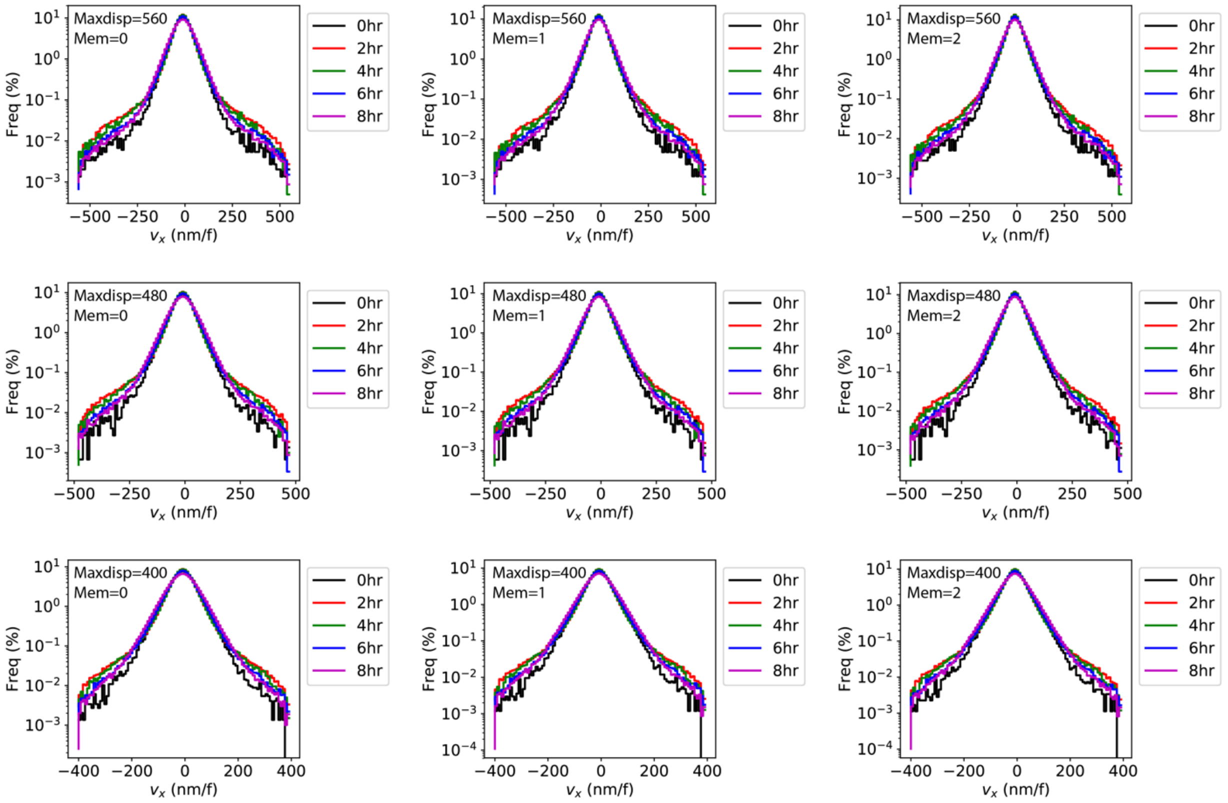


**Figure S4**. Effects of varying tracking parameters, Maxdisp and Mem, on the distribution of instantaneous velocities ($v_x$).

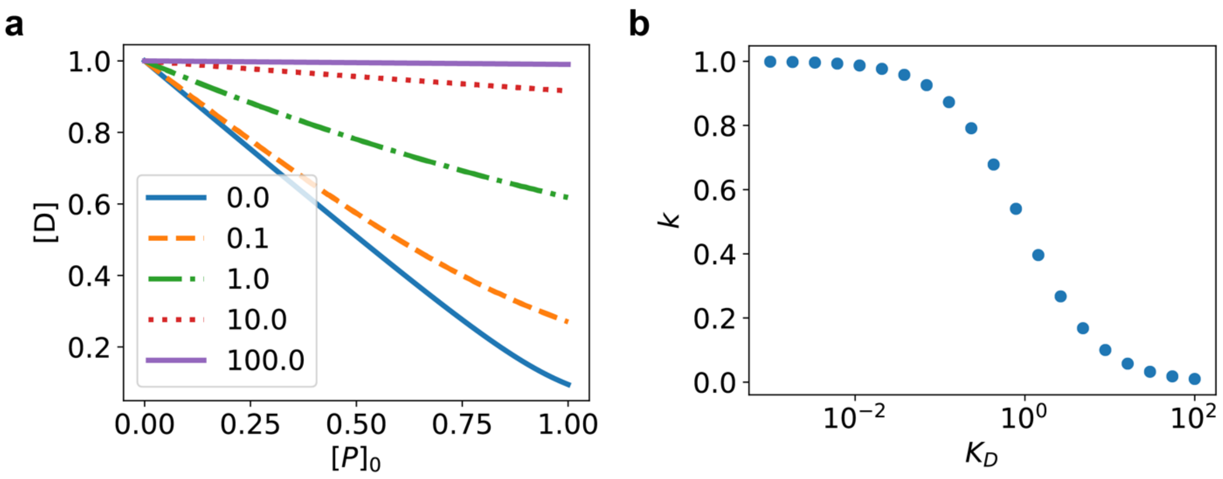


**Figure S5**. (a) Theoretical dependence of percentage of unbound DNA $[D]$ on the initial concentration of proteins $[P]_0$ with various dissociation constant $K_D$=0.01, 0.1, 1, 10, and 100, assuming $[D]_0 = 1$. (b) Computed dependence of fitted $k$ (using $[D] = [D]_0 - k \times [P]_0$) on the dissociation constant $K_D$ assuming $[D]_0 = 1$.

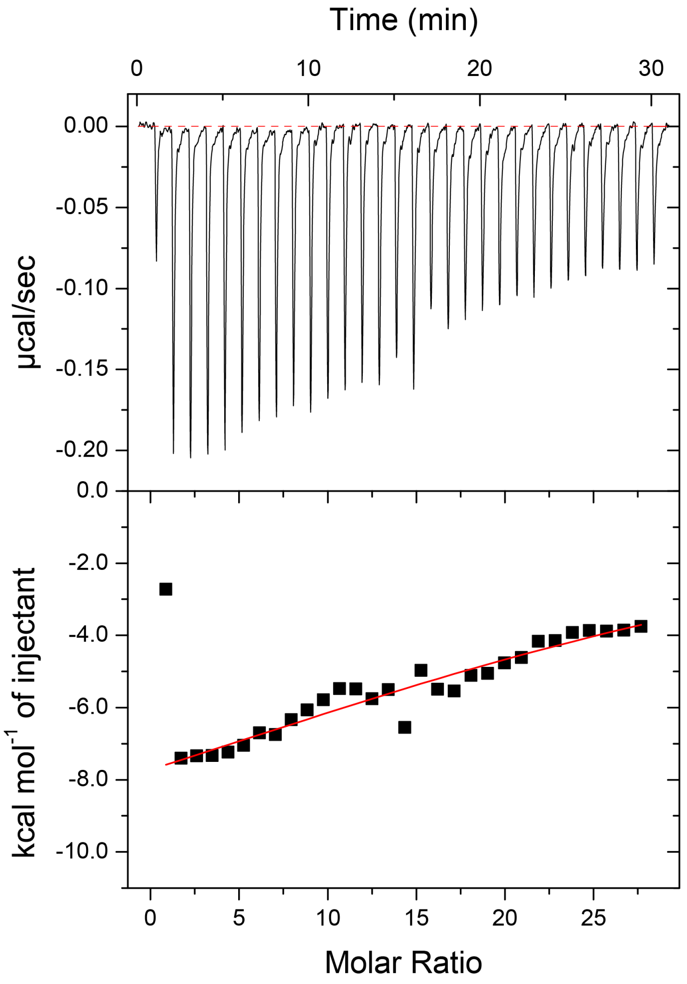


**Figure S6**. The ITC measurement of titrating 0.2 mM $Ag^{+}$ ions into DNA at pH 7.4 in 0.2 mM tris-HCl and 0.25 mM NaCl at 25 °C in 10 mM HEPES buffer. The injection volume was 1.3 µL with 1 min interval between injections. Fitting the data with the one-set binding model (red solid line) gave $K = (5.29 \pm 3.37) \times 10^{4}$ $M^{-1}$ and $\Delta H = -12.63 \pm 3.66$ kcal/mol.
